# Foxg1 and companions: not only transcription factors

**DOI:** 10.1101/2024.12.16.628377

**Authors:** Antonello Mallamaci, Osvaldo Artimagnella, Gabriele Liuzzi

## Abstract

Here, moving from our most recent results on Foxg1 biology, we firstly summarize available information about a few, special pleiotropic effectors of neurodevelopmental interest, involved in control of both transcription and post-transcriptional steps of gene expression. Next, upon further scanning of literature, we report evidence that, not strictly limited to neurodevelopmental processes, such functional pleiotropy also applies to other transcription factors, involved in physiology and homeostasis. Besides, by systematic mining of a major public protein-protein interaction database, we collect robust evidence that an involvement of “canonical” transcription factors in post-transcriptional control of gene expression may be a pervasive phenomenon, characterizing hundreds of effectors. Finally, we discuss the biological meaning of these findings and propose three evolutionary mechanisms that may have conspired to such unexpected scenario.

## MAIN TEXT

### Functional pleiotropy of Foxg1 and other transcription factors mastering key neurodevelopmental processes

Classically known as a transcription factor mastering the development of the mammalian rostral brain and promoting cytoarchitectural maturation and activity of neocortical neurons (summarized in ref^1^), Foxg1 has been more recently shown to be engaged in an array of activities different from pure transcriptional control.

Inspired by previous detection of Foxg1 in neuronal cytoplasm^2^ and published protein-protein interaction data^3^, we recently found that Foxg1 tunes translation of hundreds of genes, by modulating the recruitment of their mRNAs to holo-ribosomes as well as the progression of the latter on the former. This is associated to physical interaction between Foxg1 and specific translation factors (EIF4E and EEF1D) and, remarkably, can be replicated by an artificial, cytoplasm-confined variant of Foxg1 unable to impact transcription^1^. To note, genes undergoing translational Foxg1 control are often deeply involved in neuronal physiology. They include, for example, *Grin1*, encoding for the main subunit of the NMDA receptor^1^.

Remarkably, non-transcriptional impact of Foxg1 on gene expression is not restricted to translation, but also extends to other processes, including retro-transcription and post-transcriptional RNA processing.

In this respect, we recently found that Foxg1 modulates the activity of L1 retrotransposons, whose transcription is instrumental to sustain neocortical histogenesis^4^ and whose somatic amplification usually accompanies this process^5^. Specifically, we found that, while repressing *L1* transcription in committed neuronogenic progenitors and postmitotic neurons as a “professional transcription factor”, Foxg1 also promotes retro-transcription of the corresponding mRNA, by functionally counteracting two retro-transcription-inhibiting helicases, Mov10 and Ddx39a^6^, which physically interact with Foxg1^3^ and have their mRNAs paradoxically upregulated by it^6^. In this way, Foxg1 plays a prominent role in promoting the increase in *L1* copy number that normally characterizes the neonatal rodent neocortex^6^.

Furthermore, it has been shown that the decline of hippocampal miR200a level evoked by Foxg1 hemi-ablation exceeds by more than four-folds the parallel decline undergone by pre-miR-200a. Moreover, in neural N2a cells, Foxg1 binds to Drosha via a Ddx5 bridge, both are connected to pri-miR200b/200a/429, and Foxg1 promotes microprocessor activity^7^. All that suggests that, in addition to stimulating the synthesis of *pri-miR200b/200a/429* transcription^7^, Foxg1 is likely involved in early steps of its post-transcriptional processing to mature miRNAs^7^. To note, this may conspire to fine tuning of *Foxg1*-mRNA levels, sensitive to miR200-family effectors^8–10^, and impact other key metabolic variables, such as plasticity-related PKA activity^11^, dampened by the product of miR200-sensitive *Prkar2b*-mRNA^7,12^.

Next, we have found that Foxg1 overexpression in primary neocortical cultures exerts a profound impact on splicing and polyadenylation of pre-mRNA, resulting in >100 genes with altered splicing and >1300 ones with altered polyadenylation (ref^1^ and our unpublished data). However, we presently ignore the underlying molecular mechanisms, and we cannot rule out that such scenario may “simply” originate from altered transcription of genes encoding for key factors implicated in splicing and polyadenylation control.

Finally, in this context, it is worth mentioning the special liaison that links Foxg1 to mitochondria. Two specific isoforms of it, ^45kD^Foxg1 and ^25kD^Foxg1, likely originating from proteolysis of the major ^58kD^Foxg1 one, are localized in the inner mitochondrial matrix, whereas ^58kD^Foxg1 prevails in the nucleus and ^45kD^Foxg1 in the cytoplasm^13^. Experimental overexpression of ^58kD^Foxg1 and ^25kD^Foxg1 results in a decrease and an increase of median mitochondrial size, respectively^13^. Conversely, *FOXG1* haploinsufficiency reduces the number of mitochondria per cell and their branching index, while not affecting their average size^14^. Moreover, overexpression of ^58kD^Foxg1 and ^25kD^Foxg1 abolish the respiratory reserve, i.e. the maximal-supplementary, mitochondrial O2 consumption ability, exceeding that normally employed by these organelles to sustain ATP synthesis, while not affecting O2 consumption rates linked to ATP generation^13^. Conversely, *FOXG1* haploinsufficiency leads to lower cellular ATP content, possibly as a consequence of diminished mitochondrial mass per cell^14^. At the moment, we ignore if mitochondrial Foxg1 isoforms are implicated in these phenotypes via extra-transcriptional activities. To note, as many as two out the four FOX-family transcription factors known so far to localize in mitochondria, FOXO1 and FOXO3, bind to the D-loop^15,16^, namely a large, triple-helix- and regulatory elements-rich region of the mitochondrial chromosome, implicated in its both transcriptional and replicational control^17^.

Quite in advance to recent functional dissection of Foxg1, an involvement in post-transcriptional control of gene expression had been already described in case of a few homeodomain-TFs, prevalently involved in embryo patterning.

The very first was the Drosophila *bicoid* (*bcd*) gene, encoding for a homeodomain-TF expressed in the syncytial-blastoderm of the fly embryo along an anterior^high^-caudal^low^ gradient, and acting as a maternal determinant of anterior identity. As early as in 1996, it was reported that, in addition to differentially promote transcription of three anterior gap genes, *hb*, *otd* and *kr*^18–20^, Bicoid also trans-represses translation of the ubiquitous *cad*-mRNA, so limiting high expression of its protein product to the caudal-most part of the embryo^21^. It was found that Bicoid binds to a BRE-motif lying in *cad*-mRNA-3’UTR, by means of an arginine-rich module (ARM) located in the 3’-most part of its homeodomain^22^. Moreover, anchored to *cad*- mRNA, Bicoid further binds to EIF4E appended to *cad*-mRNA-5’cap, thanks to a YXXXXLΦ motif similar to those mediating the EIF4E/EIF4E-BP interaction. In this way, Bicoid prevents EIF4E interaction with EIF4G and therefore jeopardizes *cad*-mRNA translation^23^. To note, *bicoid* appeared in long-germ-band dipterans like Drosophila, where it arose from a duplication of the Hox3 homologue zerknullt^24^. As for *caudal*, albeit present even in short-germ-band dipterans, differently from flies, it is *dynamically* expressed by the caudal proliferating blastema of these more ancestral insects, from which all segments posterior to pre-gnatal head are progressively generated^25^. As such, i.e. because of the *variable distance* between its expression domain and the anterior pole of the embryo, short-germ-band dipteran *cad* gene might hardly be engaged in an interaction similar to the *bcd/cad* one taking place in flies. All this suggested that *bcd* control of *cad* translation might be just the result of a “fortuitous evolutionary accident” having occurred within a highly divergent phyletic lineage, such as the Drosophila’s one.

However, shortly afterwards, several studies were published, showing that it is not so and an involvement in post-transcriptional control of gene expression characterizes a small set of other homeodomain-TFs, prevalently implicated in vertebrate embryogenesis control. In 2005, it was reported that, specifically in myeloid cells, the Proline-Rich-Homeodomain transcription factor (PRH) binds to EIF4E through the YXXXXLΦ motif, known to also mediate EIF4E/EIF4E-BP1 interaction. In this way it antagonizes nuclear-EIF4E-depedent, nucleo- cytoplasmic transport of *CyclinD1*-mRNA, which reduces its translation, preventing myeloid cell transformation^26^. Within the same cells (and via the same YXXXXLΦ motif), Hoxa9 homeobox TF can bind to nuclear-EIF4E too, outcompeting PRH and so promoting nucleo- cytoplasmic transport of both *CyclinD1*- and *ODC*-mRNA. Limited to *OCD*, Hoxa9 also increases recruitment of its mRNA to polysomes, leading to a further upregulation of its post-transcriptional expression gain. All this may contribute to transformation of myeloid cells^27^. In 2005, it was also reported that an extracellular source of mesencephalon-patterning, En2 homeobox TF specifically repels and attracts axonal growth cones originating from the amphibian temporal and nasal retina, respectively, within a timeframe too short for a transcription-mediated mechanism. As expected, this phenomenon occurred even upon severing axons, was suppressed by anysomycin and rapamycin (but not by α-amanitin), and was associated to a local increase of translation rates. Intriguingly, it was preceded by a rapid surge of EIF4E and EIF4E-BP1 phosphorylation levels, likely instrumental to it^28^. Consistently, four years later, it was shown that exposure of adult mouse midbrain *synapto-neurosomes* (*not whole cells*) to the En2 paralog, En1, was followed by an *acute* selective increase of nuclearly-encoded, Nduf1 and Nduf3 mitochondrial complex I subunits, which resulted in increased complex I activity. To note, Nduf1 and Nduf3 levels were consistently reduced in *En1^-/+^;En2^+/+^* mice, more susceptible to 6-hydroxydopamine and a-synuclein-A30P toxicity^29^. Finally, strong hints to an involvement in post-transcriptional control of gene expression emerged for *Otx* and *Emx* genes, among the earliest ones mastering CNS patterning along the coordinated axes^30,31^. A localization in olfactory axon terminals, pointing to a likely implication in peripheral control of translation, was described for Emx2 and its Emx1 paralog^32,33^. In this context, Emx2 was found to bind EIF4E via a YXXXXl<λ domain, highly similar to those mediating reciprocal interactions among EIF4E and EIF4G, EIF4E-BPs, Bicoid, PRH, Hoxa9, and En2^33^. To note, like for Hoxa9 and PRH, Emx2 involvement in post-transcriptional gene control seems to be not restricted to translation, but to also apply to other steps of mRNA metabolism, as suggested by its documented physical interaction with Cnot6l an d QkI-7^34^. More recently, mass spectrometry analysis of OTX2 protein interactors in the adult retina, as well as in choroid plexus and CNS regions non cell-autonomously sensitive to Otx2 manipulations (e.g. visual cortex and subventricular zone), revealed a large set of proteins implicated in splicing, nucleo-cytoplasmic mRNA export and translation (including U2af1, U2af2, Hnrnpk, Nxf1, Pabpc1, EEF1a1), which, similarly to Emx2, points to a likely extra-transcriptional control exerted by Otx2 on these processes^35,36^.

### Is direct transcription factor involvement in post-transcriptional modulation of gene expression limited to a few neurodevelopmental effectors or is it a pervasive phenomenon?

Prompted by these findings, we wondered if, beyond homeodomain- and winged-helix-TFs implicated in neurodevelopmental processes, other TFs might straightly modulate gene expression downstream of transcription. To preliminarily address this question, we scanned literature for reports documenting a functional impact of TFs on translation, independent from transcriptional control. We found about a dozen of these reports^37–50^ (**Table 1**), which prompted us to assess if this may be a more general and widespread phenomenon.

**Table 1.**
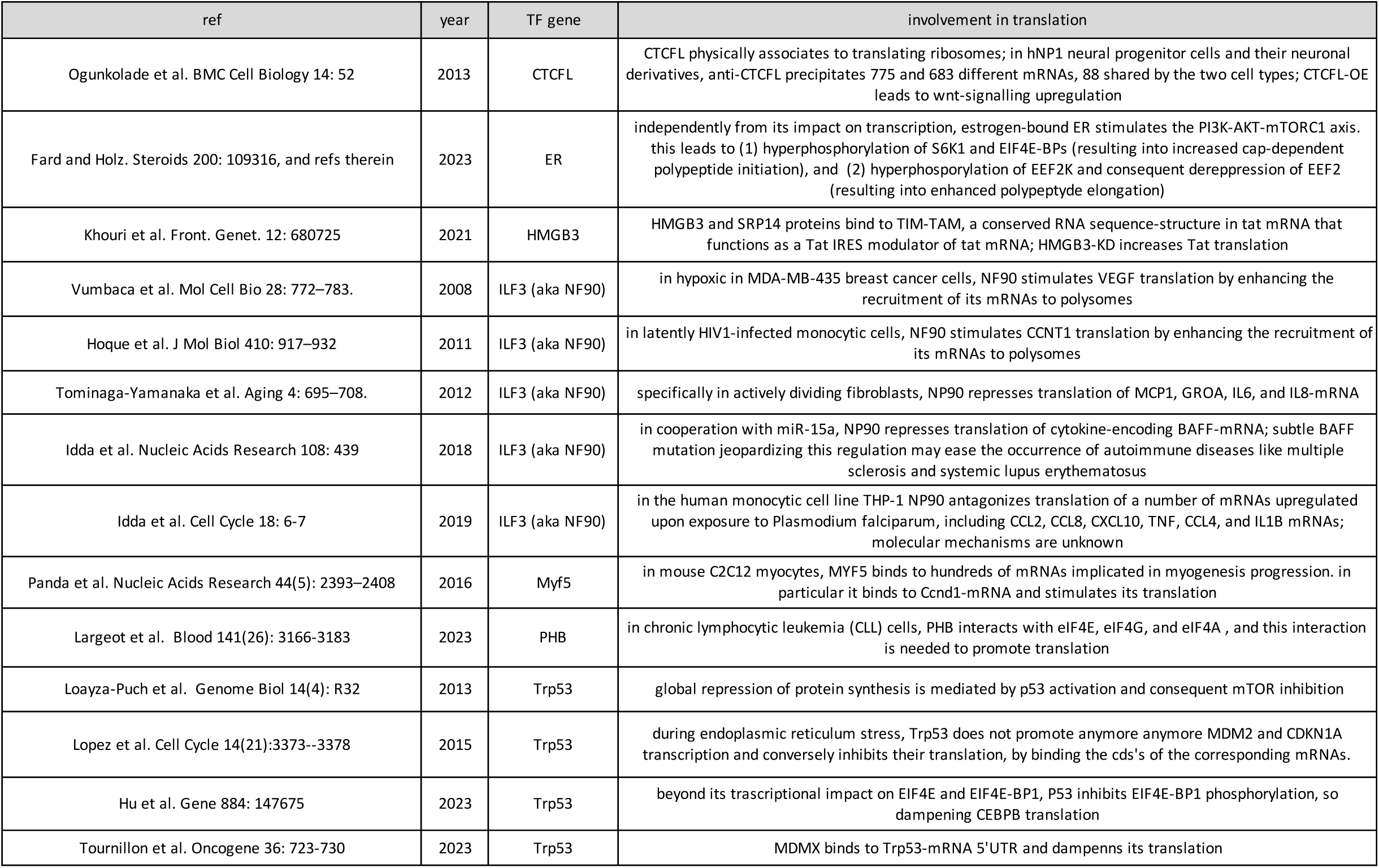
Selected publications reporting TF involvement in translational control.

To systematically address this issue, we decided to capitalize on huge protein-protein interaction data stored in public databases, considering the specific physical interaction between a given TF and classical factors controlling distinctive post-transcriptional steps of gene expression (splicing, polyadenylation, translation) as a reasonable index of possible direct control exerted by such TF over these processes. Our analysis strategy was as summarized in **Figure 2** and detailed below.

**Figure 1.**
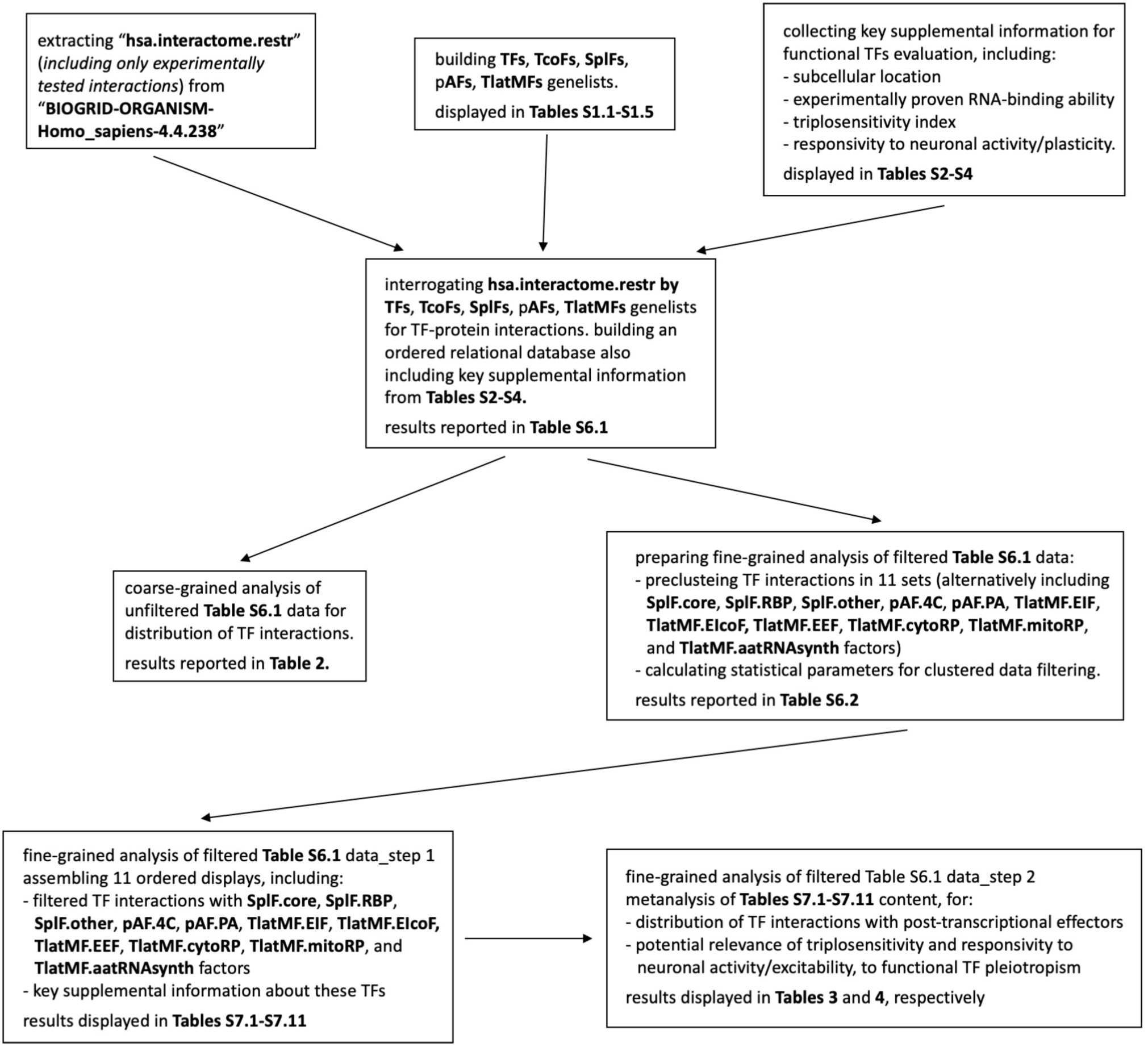
Mining the BioGrid database: operational diagram. Here shown are the structure and the concatenation of the main workpackages implemented to extract key information supporting large scale TF implication in post-transcriptional control of gene expression.

**Figure 2.**
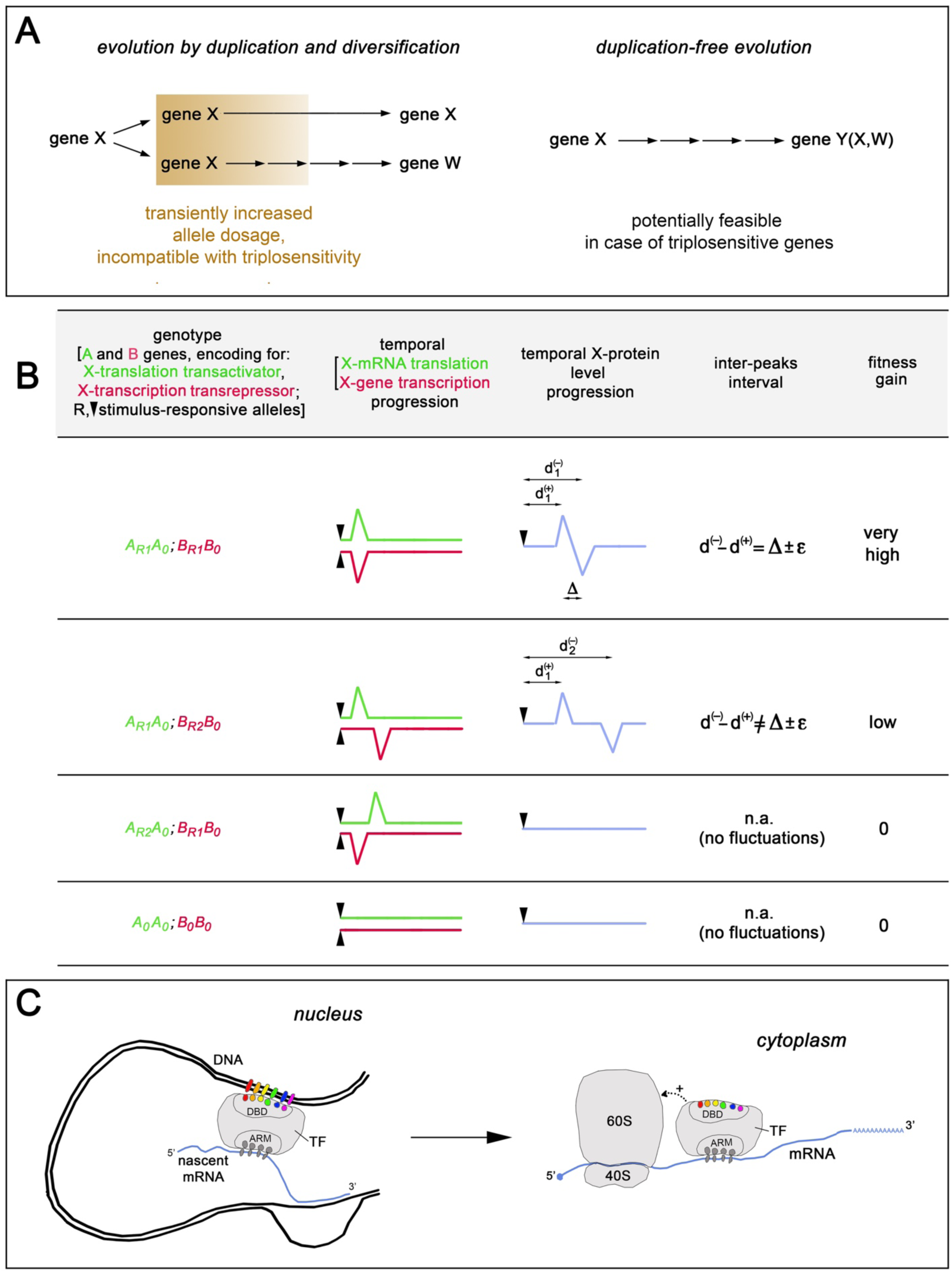
Hypothetical mechanisms conspiring to evolution of pleiotropic TF control over transcription and post-transcriptional steps of gene expression. **(A)** Implicating a transient increase of allele dosage, the generation of a novel W gene by duplication and diversification of an X ancestor would lead to a corresponding fitness drop, if the organism is X- triplosensitive. That might have pushed organisms to follow alternative, duplication-free pathways, whereby a triplosensitive X-ancestor progressively evolves into a Y derivative, encoding for a protein co-executing both X and W functions. **(B)** Timed fluctuations of translation and transcription rates of an X gene, driven by transactive products of two, A and B, stimulus-responsive genes, respectively, can lead to peculiar temporal fluctuation profiles of X protein. For example (row 1 of the prospect), transient, simultaneous transactivation and transrepression of translation and transcription, respectively, can generate two sequential X peaks, a positive and a negative one, separated by a Δ interval. In a given context, this may confer to the organism harboring *AR1* and *BR1*, stimulus-responsive alleles, needed for such response, a remarkable fitness gain, paving the way to selection of such alleles. However, the partial replacement of these alleles by other stimulus-responsive ones, characterized by different response-latencies (*AR2* and *BR2*; see rows 2 and 3 of the prospect), can reduce or also abolish this fitness gain. For this reason, the *average* fitness gain associated to *AR1* and *BR1* can be quite lower, compared to the *maximal* one that they elicit when reciprocally associated. This considerably delays their selection. Given such scenario, every device that keeps the capabilities to encode for AR1 and BR1 functions together can tremendously increase the *average* fitness gain of the corresponding individuals, speeding up its selection. Conveying transcriptional and translational control abilities on the product of the same gene can be considered just the limit strategy to achieve this goal. **(C)** Transcription factors (TF) bind to DNA motifs localized in 3D-surroundings of specific transcription units via dedicated DNA- binding domains (DBD), in a high specific fashion (interacting aa-residues and nucleotides in color). The same TFs may co-transcriptionally interact with nascent mRNA molecules, via RNA- binding domains (ARMs), supporting a less specific interaction (interacting aa-residues and nucleotides in grey), however sufficient to sustain a relatively stable association between the two molecules, which survives mRNA detachment from DNA and its nucleo-cytoplasmic translocation. In this way, recruited *specifically* at a given gene, a TFs may *selectively* modulate post-transcriptional metabolism of mRNAs originating from that gene. In other words, conveying transcriptional and translational control abilities on the product of the same gene, may allow to rapidly “recycle” the evolutionarily laborious selection of specific DNA/protein interfaces for advantageous, indirect drive of specific RNA/protein interactions.

We compiled a number of genelists (TFs, transcription factors; TcoFs, transcription cofactors; SplFs, splicing factors; pAFs, polyadenylation factors; TlatMFs, translational machinery factors; **Tables S1.1-S1.5**), and - by these genelists - we mined the BIOGRID-ORGANISM- Homo_sapiens-4.4.238 database, limiting our analysis to experimentally verified, protein-protein interactions (“hsa.interactome.restr”). We plotted the number of interaction reports involving each TF with all distinctive post-transcriptional factors mentioned above. As a control, we also included into this plot full data referring to “canonical” TF/TF and TF/TcoF interactions (**Table S6.1**).

Starting from this ordered dataset, we calculated the cumulative numbers of TF interaction reports co-including factors encoded by each of the five genelists mentioned above (TFs, TcoFs, SplFs, pAFs, TlatMFs). Moreover, we repeated such cumulative assessment limited to TFs belonging to distinctive taxa (**Table 2**). Remarkably, the total number of TF/SplF and TF/TlatMF interaction reports (13,287 and 6,240) was not negligible if compared to TF/TF and TF/TcoF ones (35,728 and 31,881). Besides, upon normalization against TF/TF and TF/TcoF values, TFs belonging to some taxa (HMG.WHSC1, NDT80, ZF.BED, ZF.C3H, ZF.CCHC, ZF.NFX1, IPT, HMG.WHSC2, ZF.CXXC, DBP, HSF, PUR, ZF.DM, WC.MYB, CSRNP_N, HMG, RUNT, HMG.TOX) resulted to interact at particularly high frequencies with SplFs, TFs belonging to other taxa (ZF.CCHC, ZF.NFX1, DBP, HMG.TFAM, HMG, PUR, GHD.CP2) with TlatMFs. To note, albeit represented in the database by several hundreds TF/TF and TF/TcoF interaction reports, some taxa (TEA, NF1, MADS, HB-CUT, ARID, HB-PAX, PAX) did not display any TF/TlatMF entries. Altogether, these results suggest that: (1) TFs might be specifically implicated in non-canonical post-transcriptional control of gene expression, to an extent sometimes comparable with their canonical involvement in transcriptional control; (2) TFs belonging to distinctive families might be differentially engaged in splicing and translation control.

**Table 2.**
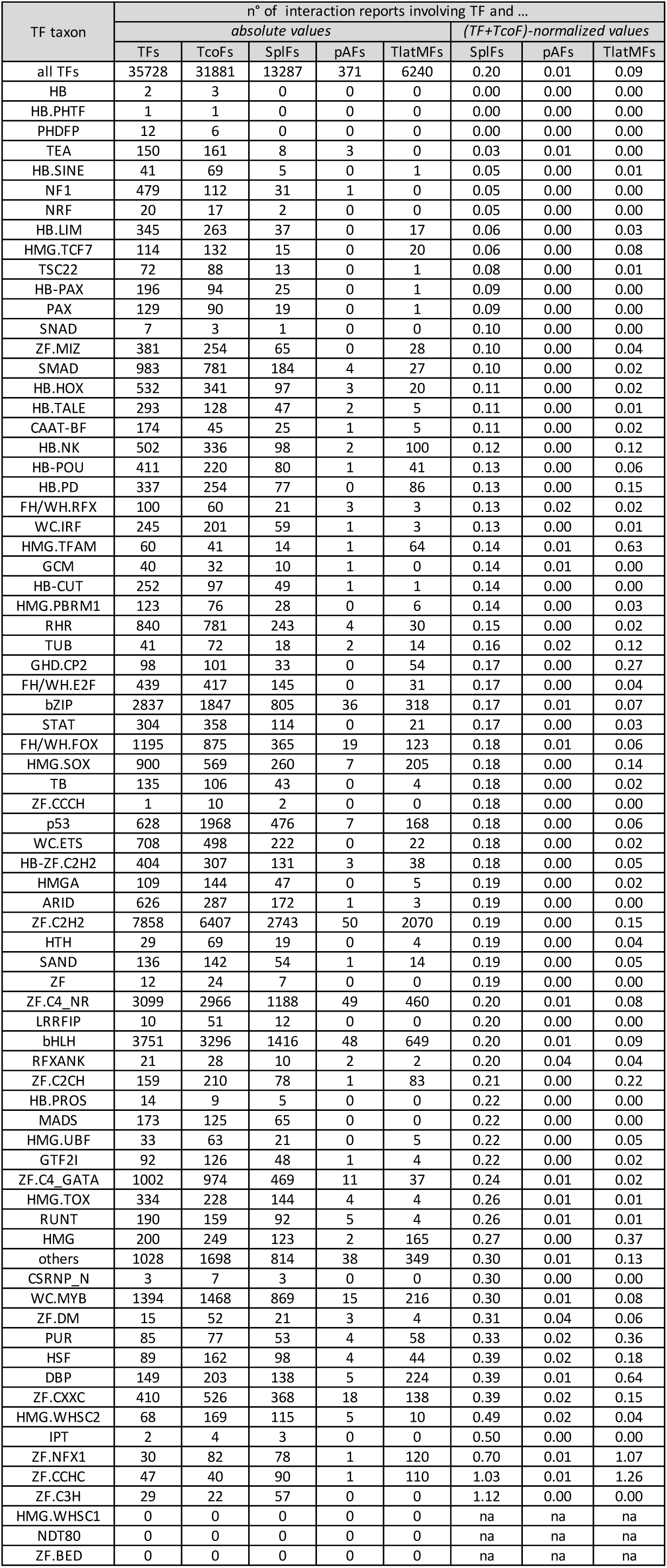
Coarse-grained distribution of full BioGrid, TF-interaction reports (1).

Surprised by this scenario, we suspected that it could have trivially stemmed from aspecific TF interactions, occurring upon protein overexpression needed for the interaction assays indexed in BioGrid. If so - we reasoned - frequencies of BioGrid interaction reports should give rise to a pattern not far from that originating from random proteins inclusion into interactor dyads, reflecting their cumulative representation frequencies in the database. Therefore, we decided to challenge our original inference based on *coarse-grained* analysis of the *full* dataset, by (1) analyzing a *more finely-grained* dataset generated from **Table S6.1** data upon clustering interaction reports involving each TFs and 11 smaller distinctive sets of functionally related post-transcriptional effectors (SplF.core, SplF.RBP, SplF.oth(3), pAF.4C, pAF.PA, TlatMF.EIF, TlatMF.EIcoF, TlatMF.EEF, TlatMF.cytoRP, TlatMF.mitoRP, TlatMF.aatRNAsynth; see **Tables S1.1-S1.5**), and (2) *retaining only those interaction-report sets* displaying a statistically significant excess of observed over random-expected items (**Table S6.2**). In this way, we ended up with 11 tables, where we synthesized “filtered” experimental evidence in support of TF implication in splicing (**Tables S7.1-S7.3**), polyadenylation (**Tables S7.4,S7.5**) and translation (**Table S7.6-S7.11**). We further summarized key features of this Table-set in **Table 3**, where - for clarity of narration - TFs were displayed as “preys” captured by pools of “baits” associated to distinctive aspects of splicing, polyadenylation and translation.

**Table 3.**
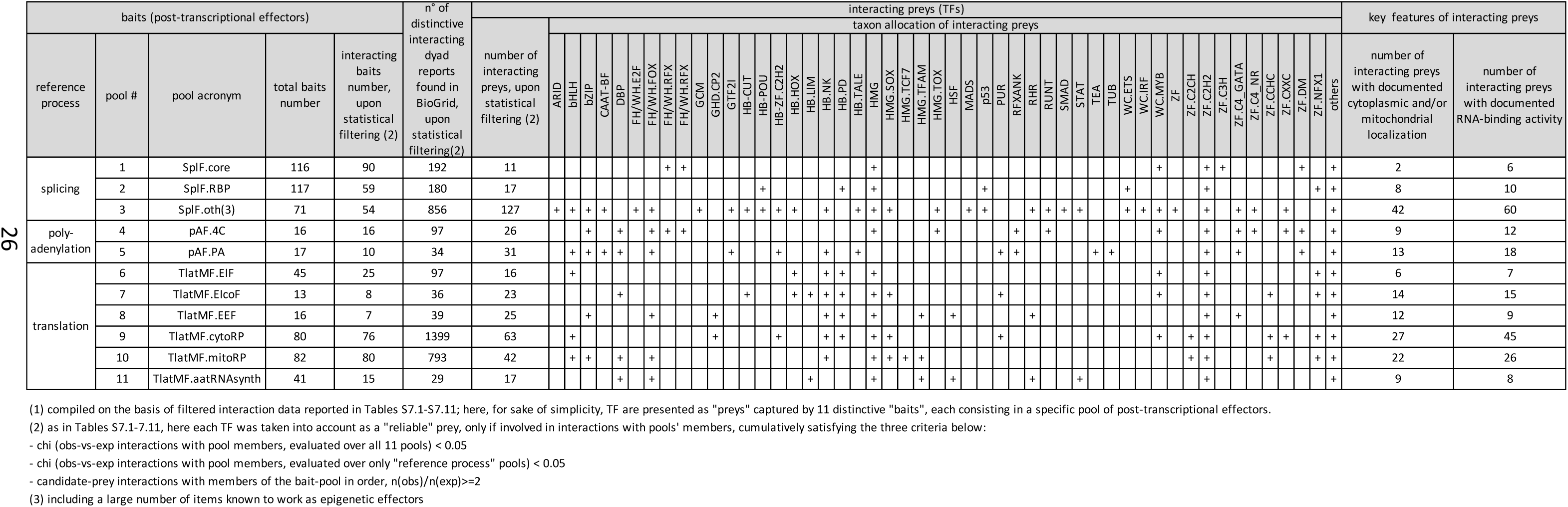
Distribution of physical interactions among classified TFs and post-transcriptional factors, upon statistical filtering of primary BioGrid data (1), with selected key features of interacting TFs.

To note, where applicable, for each TF, we also annotated (1) its extranuclear localization (as documented in the Human Protein Atlas (**Table S2**)), and (2) its experimentally verified capability to bind RNA (according to a full list of human RNA-binding proteins, RBPs, precompiled by us on the basis of primary literature (**Table S3**)), namely two valuable pieces of information, potentially suitable to corroborate inferred TF control over specific post-transcriptional steps of gene expression. To ease subsequent correlative analyses, these data were included in **Table S6.1** and its **Table S7.1-S7.11** derivatives.

As shown in **Table 3**, as many as 398 TFs, i.e. about 25% of those subject of investigation, specifically interact with post-trascriptional controllers of gene expression and, as such, are likely involved in transcription-independent modulation of these processes. These TFs include 155 ones involved in splicing, 57 in polyadenylation, and 186 in translation, cumulatively interacting with 203, 26 and 211 post-trascriptional factors canonically implicated in these processes, respectively.

Remarkably, beyond the stringent data filtering procedure mentioned above, as many as four specific a posteriori arguments (see **Table 3**) suggest that such pattern emerged from genuine interactions among these effectors and does not trivially reflect random, aspecific contacts among them. First, *very different fractions* of post-transcriptional effectors included in distinctive bait-pools are engaged in interactions with TFs, from as little as 36.6% of aa.tRNA.synthetases to 100% of pAF.4C members. Second, the correlation index between the number of bait-pool members engaged in interactions with TFs and the number of their preys is very low, close to 0.26. Third, the allocation of preys captured by distinctive bait-pools to different TF taxa is robustly diversified, being the average value of such diversification close to 33%. [To note, such diversification is remarkably pronounced in case of TlatMF.cytoRPs and TlatMF.mitoRPs pools (**Tables 2, S7.9** and **S7.10**), which, together with high TF-connectivity of TlatMF.mitoRPs, rules out that high TF-connectivity to TlatMF.cytoRPs may trivially reflect geometrical TF-proteins contiguity to ribosomes at the time of their synthesis]. Finally, inspecting our lists of TFs putatively engaged in translational control (**Tables S7.6-S7.11**) revealed that they collectively included only three TFs previously proven to be implicated in this control (EMX2, FOXG1 and PHB, see **Table 1**), while missing the remaining six (CTCFL, ER, HMGB3, ILF3, MYF5, EN2 and OTX2). This suggests that our statistical filtering strategy was likely too severe, leading us to underestimate the pervasivity of phenomena subject of investigation (in this respect, compare **Table S6.1** with **Tables S7.1-S7.11** contents).

Interestingly, for a substantial fraction of TFs interacting with translational machinery (90 out of 186), an extranuclear subcellular localization is reported (**Tables 3** and **S2**). Moreover, for >1/2 of TFs putatively involved in non-canonical control of post-transcriptional gene tuning (76/155, 30/55 and 110/188, in case of splicing, polyadenylation and translation, respectively), the capability to bind RNA in addition to DNA has been experimentally verified (**Tables 3** and **S3**). This suggests that, in principle, these TFs might modulate post-transcriptional steps of gene expression in a gene-specific way, bridging canonical splicing-, polyadenylation- and translation-factors to specific mRNAs or - alternatively - inhibiting their reciprocal interaction.

In sum, based on protein-protein and protein-RNA interaction data available from public databases, we predict that (1) in addition to canonical transcription control, at least 1 out 4 of our TFs can be non-canonically implicated in control of post-transcriptional steps of gene expression, with a particular emphasis on translation, and, (2) in up to one half of these cases such non-canonical control might be exerted in a gene-specific way.

Albeit based on a *conservative* statistical evaluation of a huge body of interaction data, this prediction obviously requires caution. In fact, a large subset of TF interactions on which it is based might simply represent metabolic noise. Alternatively, they might be pertinent to processes different from post-transcriptional gene tuning. For example, massive TF “sequestering” by ribosomal proteins (>2100 filtered, distinctive interacting dyads found, **Table 3**) might be instrumental in dampening canonical TF activity, by tightening TF confinement to extranuclear compartments. On another side, the widespread TF ability to bind RNA might mainly reflect the emerging, pervasive employment of RNA cofactors to support specific aspects of transcriptional mechanics (reviewed in ref^51^). Consequently, our prediction does require *functional* experimental validation. In this respect, CASH^52^ and ROAR^53^ (re-)analysis of RNASeq data from TF-OE and/or TF-LOF preparations could help probing candidate TF involvement in splicing and polyadenylation, respectively. Similarly, mild overexpression of *cytoplasm-confined* TF-chimeras^1^ followed by delayed proteome profiling might make *serial* assessment of straight TF implication in translational control a feasible task.

### Why transcription factors pleiotropy?

Beyond the experimental validation issues mentioned above, the pervasive funneling of supplemental control functions on molecular effectors already engaged in transcriptional control is theoretically puzzling. It is commonly accepted that, during evolution, genes encoding for polypeptides endowed with new functions generally arose by duplication and diversification of pre-existing genes^54–57^. In fact, employing just an essential gene for evolutionary experimentations could pose obvious risks for the fitness of a species. Convesely, employing duplicates of it would allow to circumvent this problem, via the generation of new “semifinished substrates” freely diversifiable to encode novel functions, thanks to the release of the pressure acting on such essential gene. If so, why as many as one quarter of our TF genes (or more) would encode for (sophisticated) multi-task effectors? We hypothesize that three distinctive evolutionary factors may have conspired to such scenario: (1) the triplosensitivity bottleneck (2) the possibility to optimize the inheritability of complex spatio-temporal gene regulation patterns (3) the possibility to achieve fast evolutionary transfer of pre-selected, protein-DNA interaction specificities to protein-RNA interactions. Below, we detail these hypotheses and discuss how assessing their goodness.

Concerning the first hypothesis, it has been shown that proper allele dosage is often crucial to individual health. In particular, ≈3,000 of our autosomal, polypeptide-encoding genes have been inferred to be haploinsufficient, and ≈1,000 of them also triplosensitive^58^, with an obvious link between anomalous CNVs and impaired mental health^59^. On the other hand, it has been estimated that, albeit fast, structural gene diversification which follows gene duplication, requires a *not-negligible* evolutionary time^60^. Because of that, evolution of triplosensitive genes can hardly exploit the “duplication & diversification strategy”. Such strategy - in fact - would expose individuals adopting it to an enduring fitness drop. That is why these individuals are “pushed” towards an alternative path, i.e., funneling novel functions on the protein product of their *original* gene (**Fig. 2A**).

To probe the goodness of this inference, we took advantage of “pTriplo”, namely a numerical index originating from accurate meta-analysis and machine-learning exploitation of clinical data from almost 1,000,000 individuals, which provides a reliable triplosensitivity prediction for each autosomal gene^58^ (**Table 3**). We found that, while equalling 0.19 among all autosomal genes, the fraction of genes with pTriplo≥0.75 arose to 0.33 among all TFs and further to 0.47 among TFs engaged in interactions with post-transcriptional effectors (“pt-TFs”, with observed-interaction-freq>3*expected-interaction-freq, and p<0.001). Besides, more and more pronounced increases characterized the fractions of genes with progressively higher pTriplo. Specifically for pTriplo ≥0.99, such fraction equalled 0.03, 0.06, 0.09, and 0.11, among total genes, autosomal TFs, pt-TFs with observed-interaction-freq>2*expected-interaction- freq and p<0.05, and pt-TFs with observed-interaction-freq>3*expected-interaction-freq and p<0.001, respectively. All that corroborates the hypothesis that triplosensitivity may be a major driver of gene duplication-free evolutionary pathways, such as those proposed for TFs involved in post-transcriptional control of gene expression.

Concerning the second hypothesis, optimal unfolding of information-rich metabolic processes (with a special emphasis on those implemented by CNS and/or immune cells) may often require a temporally structured, often non-monotonic, regulation of levels of gene products involved in their implementation. Of course, this requirement can be easily fulfilled by orthogonal tuning of two distinctive steps of gene expression (e.g., transcription and translation), exerted by two dedicated, A and B effectors, and can confer an appreciable evolutionary advantage to the individual able to perform it. Such advantage, however, only emerges, provided that the temporal activity profiles dictated by the alleles encoding for these effectors are properly coupled. In other words, type-X allele encoding for effector A elicits an appreciable fitness-gain only if working in concert with type-Y allele encoding for effector B (**Fig. 2B**). It turns out that each device increasing the linkage between AX and BY alleles can increase the *average* fitness-gain associated to each of them, so speeding up their positive selection. In this context, the duplication-free evolution of an original A gene to a new A/B identity might have been implemented just as the limit-strategy suitable to achieve this goal.

To probe the goodness of this inference, we focused on a selection of genes activated according to temporally structured patterns, in vitro and/or in vivo, in response to neuronal activity^61^, and compared their prevalence among all TFs and TFs specifically engaged in interactions with post-transcriptional effectors (**Table 3**). We found that, equaling 0.16 in the former, such prevalence arose up to 0.23 in the latter. This suggests that the advantages provided by some specific, temporally structured polypeptide expression profiles may have contributed to “push” some TF genes to evolve in a duplication-free fashion.

Concerning the third hypothesis, a few preliminary considerations are in order. First, DNA-binding specificities displayed by TFs are often very high. A preferentially-bound DNA-sequence motif has been experimentally determined (or inferred from a tightly-related homolog) for Δ1,200 out of Δ1,600 human TFs^62^, and the binding preference displayed by TFs towards such motifs may exceed by ≥1,000-folds those exhibited towards other motifs^63^. RNA-binding proteins can distinguish different RNA molecules based on their sequences too, largely thanks to a variety of dedicated RNA-binding domains^64–66^. However, TFs rarely harbor canonical RBDs. Conversely, as many as 80% of them are provided of short, basic-aa-rich domains similar to the RNA-binding domain of the HIV Tat transactivator and generally proximal to DBDs (termed ARM-like domains), shown to be necessary and sufficient to mediate RNA-binding properties displayed by these TFs^67^, however hardly sufficient - because of their relatively poor complexity - to confer to a given TF the capability to discriminate among different mRNAs. Intriguingly, it has shown that a number of yeast effectors involved in fine tuning of transcription (including Rbp4, Rbpp7, CCR4-Not, Xnr) are co-transcriptionally loaded on nascent pre-mRNAs and remain specifically bound to their mRNA derivatives even upon their nucleo-cytoplasmic translocation, so modulating late post-transcriptional steps of gene expression, including mRNA translation and decay^68–75^. In other words, polypeptide effectors, previously recruited to specific DNA regions, can be *specifically* transferred to RNA emerging from transcription of these regions, likely thanks to their *transient vicinity*, and remain attached to these RNAs for a time sufficient to heavily impact their metabolism. It is tempting to speculate that conveying post-transcriptional functions on the very same gene products already involved in transcriptional control was just a smart evolutionary “trick” that made sequence-binding specificities laboriously pre-distilled against distinctive DNA sequences easily and specifically portable to mRNA products of their transcription, where they were “rapidly recycled” to implement more sophisticated regulatory programs (**Fig. 2C**). This inference obviously requires a rigorous experimental validation. In this respect, a systematic inspection of TF phylogenetic trees for possible late appearance of ARM-like domains, as well as an assessment of the *necessity of nuclear co-transcriptional loading* of TFs on pre-mRNAs to achieve their further extra-nuclear control, might be of help.

## Conclusions

Here, moving from some “unorthodox” findings reported in specialized neurodevelopmental literature, we firstly show that a pleiotropic TFs involvement in translation control had been also previously documented in other cases, not limited to the neurodevelopmental field.

Next, upon structured interrogation of a major, public protein-protein database, we report that at least one quarter of TFs, preferentially belonging to defined taxa, specifically interact with effectors implicated in splicing, polyadenylation and translation, pointing to a likely large scale, functional involvement of them in post-transcriptional control of gene expression. Consistently, we show that about one half of TFs interacting with translation machinery factors are detectable outside of the nucleus. Besides, we also show that one half of TFs putatively implicated in post-transcriptional gene control can bind RNA, meaning that their post-transcriptional control over gene expression might be implemented in an mRNA-specific fashion. Finally, we propose some experimental approaches for systematic validation of these inferences.

If confirmed, such prominent pleiotropy is theoretically puzzling, as clashing with the standard model of gene evolution by duplication and diversification. In this respect, we propose three evolutionary mechanisms which might have led to high prevalence of pleiotropy among TFs (triplosensitivity bottleneck, discrete heritability of temporal expression patterns, DNA-dependent specificity of protein-RNA interactions), and provide experimental evidence and methodological hints for validation of these proposals.

## ABBREVIATIONS LIST

*ARM*: arginine-rich motif
*DBD*: DNA-binding domain
*hsa-TF*: Homo sapiens transcription factor
*hsa-TcoF*: Homo sapiens transcription cofactor
*hsa-SplF*: Homo sapiens splicing factor
*hsa-pAF*: Homo sapiens polyadenylation factor
*hsa-TlatMF*: Homo sapiens translation machinery factor
*LOF*: loss-of-function
*OE*: overexpressing
*pAF.4C*: (human) polyadenylation factors, “4C” set (including CFIm, CFIIm, CPSF, and CSTF elements)
*pAF.PA*: (human) polyadenylation factors, “PA” set (including polyadenylate-interacting elements)
*pTriplo*: triplosensitivity index
*pt-TFs*: post-transcriptionally engaged TFs
*RBP*: RNA-binding protein
*SplF.core*: (human) splicing factors, “core” set (including U1snRNP, SM proteins, U2 snRNP proteins and related ones, A complex, LSm proteins, U5 SNP, U4/U6 SNP, PPR19 complex, B complex, Bact complex, RES complex, and C complex elements)
*SplF.other*: (human) splicing factors, “other” set (including protein kinase, Other SAPs, Minor, Lysine transferases, chromatin-related, Histone transferases, SWI/SNF Complex Components, and Polycomb Group Genes elements)
*SplF.RBP*: (human) splicing factors, “RBP” set (including (EJC/mRNP, TREX, helicases, hnRNP, SR proteins, RNA binding proteins, RNA editing, RNA modifying, RNA methylation, CBC, 3’end, RNA degradation, and RISC elements)
*TlatMF.aatRNAsynth*: (human) translation machinery factors, “aminoacyl-tRNA-synthetases” set
*TlatMF.cytoRP*: (human) translation machinery factors, “cytoplasmic ribosomal proteins” set
*TlatMF.EEF*: (human) translation machinery factors, “eukaryotic elogation factors” set
*TlatMF.EIcoF*: (human) translation machinery factors, “eukaryotic initiation cofactors” set
*TlatMF.EIF*: (human) translation machinery factors, “eukaryotic initiation factors” set
*TlatMF.mitoRP*: translation machinery factors, “mitochondrial ribosomal proteins” set

## METHODS

### Preparation of hsa.interactome.restr

The “BIOGRID-ORGANISM-4.4.238.mitab.zip” file was downloaded from the www.biogrid.org website on 2024.10.06 and the “BIOGRID-ORGANISM-Homo_sapiens-4.4.238.mitab”file was extracted from it by Unarchive software. The latter was converted into a .txt file and further processed by TextEdit and Excel software.

Originally containing 1,251,627 rows (1 row = 1 protein/protein interaction report), BIOGRID-ORGANISM-Homo_sapiens-4.4.238.mitab” was trimmed to its “hsa.interactome.restr” derivative (including 1,152,045 rows), by retaining only the interactions labeled - within the “Interaction Type” column - by the “MI:0407 (direct interaction)“ or “MI:0915 (physical association)” tags. These represent the large majority of the experimentally verified physical interactions available in the original database.

**Interrogation of hsa.interactome.restr and results presentation in Table S6.1** hsa.interactome.restr was interrogated by the five gene lists reported in **Tables S1-S5** (TFs, TcoFs, SplFs, pAFs, TlatMFs), For each gene, (1) the total number of interaction reports involving its protein product, as well as (2) the number of interaction reports involving such product and - specifically - each TF, were counted. The results of such interrogations were reported in the grids at columns G-DSJ of **Table S6.1.**

To ease later formulation of correlative inferences, four additional key pieces of information pertinent to each TF, namely subcellular location, possible RBP function, pTriplo index and responsiveness to neuronal activity/plasticity were added to **Table S6.1**, at columns DSL-DSO. They were taken from **Tables S2-S4**, previously compiled as detailed in the corresponding notes.

### Analysis of Table S6.1 data

Interaction reports data presented in **Table S6.1** were analyzed as follows.

First, all interaction reports involving (1) each single TF and (2) proteins encoded by each of the TFs, TcoFs, SplFs, pAFs, TlatMFs genesets were summarized in a grid, at columns E-I of **Table S6.2**. Information reported in this grid was then employed to compile **Table 2**.

Second, interaction reports involving (1) each single TF and (2) post-transcriptional effectors of the SplFs, pAFs, TlatMFs lists, were further analyzed for their more fine-grained topological distribution, as shown at columns K-AX of **Table S6.2**. Specifically, reports including each TF and factors implicated in post-transcriptional gene regulation were partitioned into 11 groups, corresponding to the post-transcriptional factor ensembles defined in Tables S1.3- S1.5 (SplF.core, SplF.RBP, SplF.other, pAF.4C, pAF.PA, TlatMF.EIF, TlatMF.EIcoF, TlatMF.EEF, TlatMF.cytoRP, TlatMF.mitoRP, and TlatMF.aatRNAsynth), and their actual numbers [*real values*] were reported in the grid at columns K-U. Next, a sister grid was generated, including the theoretical numbers of these reports expectable upon random interaction between the corresponding dyad members [*expected values*]. These were calculated by on basis of the frequencies of *all* reports including dyad members, evaluated over the entire database, as well as the absolute size of the database. The resulting grid is reported at columns W-AG. Then, starting from these two sister grids, three indexes-sets were calculated for each TF: (1) the “real value/expected value” ratios-set, referring to each of the 11 groups mentioned above (reported in columns AI-AS) and the “chi-square *p* values-set”, including (2) values calculated across all 11 groups or (3) limited to the 3, 2 or 6 groups forming the SplF, pAF and TlatMF ensembles, respectively (reported in columns AU-AX). The the statistical parameters mentioned above were subsequently employed to filter **Table S6.1** interaction reports included in **Tables S7.1-S7.11.** Specifically, filtered reports had to satisfy the three criteria: [a] p-chiTF (obs-vs-exp interactions with interactor group members, evaluated over all 11 groups) < 0.05; [b] p-chiTF (obs-vs-exp interactions with interactor group members, evaluated over only “reference process” groups) < 0.05; [c] reported TF interactions with members of the reference process group, n(obs)/n(exp)>=2. Finally, information reported in **Tables S7.1-S7.11** was used as substrate of the analyses reported in **Tables 3 and 4** (performed as detailed in the corresponding notes).

**Table 4.**
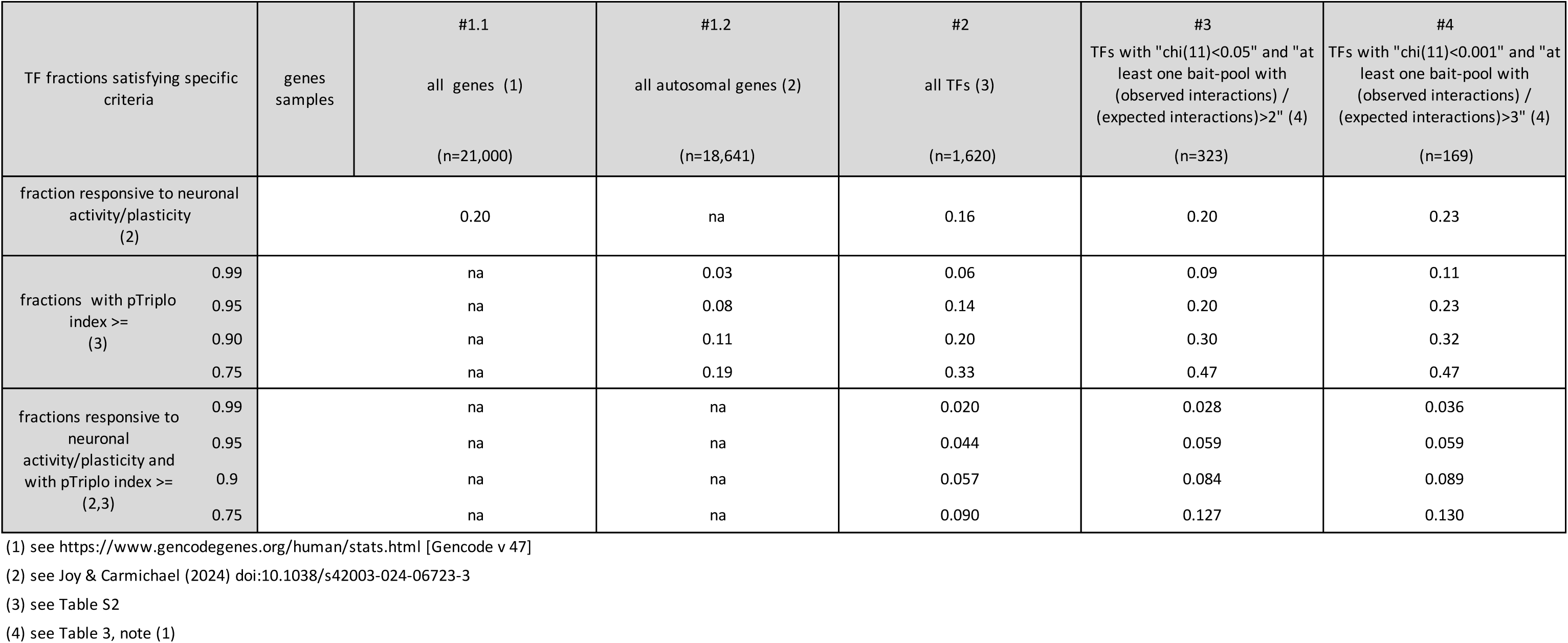
Incidence of triplosensitivity and responsivity to neuronal activity/plasticity among TFs putatively implicated in non-canonical, post-transcriptional gene control.

## AUTHORS’ CONTRIBUTION

AM conceived the study, analyzed interaction data, built Tables and Figures, and wrote the manuscript; OA took care of preliminary processing of BioGrid files; OA and GL contributed to build the RBP gene list. All the Authors discussed the preliminary version of the manuscript, provided ideas for its improvement, and read and approved its final version.

## ACKNOWLEDGEMENTS AND FUNDING

SISSA (intramural funding to AM)

1. International FOXG1 Research Foundation (Grant to AM)
2. Italian Ministery of University and Research (Grant PRIN22 2022M95RC7 to AM)
3. Fondazione Telethon (Grant GMR22T2018 to AM)

## SUPPLEMENTARY TABLES

*accessible at* https://doi.org/10.5281/zenodo.14443212

Table S1.1. hsa-TF genes list (n=1620)

Table S1.2. hsa-TcoF genes list (n=958)

Table S1.3. hsa-SplF genes list (n=304)

Table S1.4. hsa-pAF genes list (n=33)

Table S1.5. hsa-TlatMF genes list (n=275)

Table S2. Subcellular hsa-TF localization

Table S3. RBP genes list (n=4775)

Table S4. Distribution of triplosensitivity indices

Table S5. Gene relatedness to neuronal activity/plasticity

Table S6.1. TFs: full interaction data grid and other key information

Table S6.2. Full TF interact grid_statistical analysis

Table S7.1. Filtered TF/SplF.core interaction reports & related info

Table S7.2. Filtered TF/SplF.RBP interaction reports & related info

Table S7.3. Filtered TF/SplF.other interaction reports & related info

Table S7.4. Filtered TF/pAF.4C interaction reports & related info

Table S7.5. Filtered TF/pAF.PA interaction reports & related info

Table S7.6. Filtered TF/TlatMF.EIF interaction reports & related info

Table S7.7. Filtered TF/TlatMF.EIcoF interaction reports & related info

Table S7.8. Filtered TF/TlatMF.EEF interaction reports & related info

Table S7.9. Filtered TF/TlatMF.cytoRP interaction reports & related info

Table S7.10. Filtered TF/TlatMF.mitoRP interaction reports & related info

Table S7.11. Filtered TF/TlatMF.aatRNAsynth interaction reports & related info

